# Mechanisms of axoneme and centriole elimination in *Naegleria gruberi*

**DOI:** 10.1101/2024.05.22.595302

**Authors:** Alexander Woglar, Coralie Busso, Gabriela Garcia-Rodriguez, Friso Douma, Mary-Claude Croisier-Coeytaux, Graham Knott, Piere Gönczy

## Abstract

The excavate *Naegleria gruberi,* a basal eukaryote related to the “brain eating” *Naegleria fowleri*, can transform transiently from an amoeboid life form lacking flagella and centrioles to a flagellate life form where these elements are present, followed by reversion to an amoeboid state. The mechanisms imparting elimination of axonemes and centrioles during this reversion process are not known. Here, we uncover that flagella primarily fold onto the cell surface and fuse within milliseconds with the plasma membrane. Once internalized, axonemes are severed by Spastin into equally-sized fragments, which are then enclosed by membranes, after which their contents are eliminated through the lysosomal pathway. Moreover, we discovered that centrioles undergo progressive K63 autophagy-linked poly-ubiquitination and K48 proteasome-promoting poly-ubiquitination, and that ubiquitination occurs next to centriolar microtubules. Most centrioles are eliminated in lysosomes or the cytoplasm in a lysosomal- and proteasome-dependent manner. Strikingly, we uncover in addition that centrioles can be shed in the extracellular milieu and taken up by other cells. Collectively, these findings reveal fundamental mechanisms governing the elimination of essential cellular constituents in *Naegleria* that may operate broadly in eukaryotic systems.

**IMPORTANT:**

- Manuscripts submitted to Review Commons are peer reviewed in a journal-agnostic way.
- Upon transfer of the peer reviewed preprint to a journal, the referee reports will be available in full to the handling editor.
- The identity of the referees will NOT be communicated to the authors unless the reviewers choose to sign their report.
- The identity of the referee will be confidentially disclosed to any affiliate journals to which the manuscript is transferred.

**GUIDELINES:**

- For reviewers: https://www.reviewcommons.org/reviewers
- For authors: https://www.reviewcommons.org/authors

**CONTACT:** The Review Commons office can be contacted directly at

## Introduction

Flagella and centrioles play indispensable roles across eukaryotic systems. Flagella, which are also known as motile cilia, are instrumental for cell motility, whereas centrioles are crucial organizers of the microtubule cytoskeleton, being pivotal notably for cell division in animal cells. Centrioles exhibit a near-universal 9-fold radially symmetrical arrangement of microtubule blades that is imparted onto the axoneme of the flagella they template. Whereas there has been substantial progress in recent years in understanding how flagella and centrioles assemble, knowledge regarding the mechanisms governing their elimination is limited in comparison.

The removal of flagella is critical in various physiological settings and can occur through shedding into the external milieu, progressive shortening, or internalization into the cell body (reviewed in Bloodgood, 1974). In some systems, more than one mode of flagellar removal can operate, as exemplified in *Chlamydomonas reinhardtii*. Here, progressive shortening of the flagellum is catalysed by the microtubule depolymerizing enzyme Kinesin-13 (Piao et al., 2009). In addition, the microtubule-severing enzyme Katanin, which is present at the proximal end of the axoneme, enables local polymer cutting and shedding of the remainder of the flagellum (Lohret et al., 1998; Rasi et al., 2009). Observations in many other unicellular organisms established that there are distinct modes of axonemal internalization into the cell body (reviewed in Bloodgood, 1974). First, the axoneme, but not the surrounding membrane, can be retracted into the cytoplasm, as observed for example in the oral apparatus of *Tetrahymena pyriformis* (Moore, 1972; Williams and Frankel, 1973) or the zygote of *Ulva mutabilis* (Braten, 1971). Second, the flagellum can surround the cell body, after which the flagellar membrane is hypothesized to fuse laterally with the plasma membrane, resulting in axoneme internalization. This mechanism is encountered for instance also in *Ulva mutabilis* (Braten, 1971), as well as in chytrid fungi (Venard et al., 2020). In a third mode of internalization observed for example in *Allomyces arbuscula* (Barrett, 1912; Ritchie, 1947) and *Siphonaria variabilis* (Sparrow, 1937), the flagellar membrane forms a vesicle into which the axoneme rolls up and which then fuses with the plasma membrane, resulting in axoneme internalization. In the few systems where this question has been studied (e.g.: Moore, 1972; Venard et al., 2020), the axoneme is degraded following internalization through mechanisms that remain to be uncovered, although it is clear that the proteasome contributes to this process in chytrid fungi (Venard et al., 2020).

Another setting in which the fate of the flagellar axoneme can be important is after fertilization in metazoan organisms. In some systems, including mouse embryos, the flagellar axoneme is dismantled rapidly in the zygote after fertilization, such that merely axonemal fragments are present by the time of pronuclear fusion (Thompson et al., 1974). In bovine embryos, the proteasome localizes to the internalized axoneme, such that conceivably the proteasome may be involved in axoneme elimination as in chytrids (reviewed in Sutovsky, 2018). Cytoplasmic degradation of the sperm tail axoneme in the zygote could also be required to ensure that mitochondria are only contributed maternally and not from the paternal pool that resides at the sperm tail axoneme. In other systems, by contrast, the sperm-derived axoneme persists longer after fertilization. This is the case in *Drosophila* for instance, where the axoneme remains present throughout embryogenesis, with constituent axonemal proteins gradually diffusing into the cytoplasm (Pitnick and Karr, 1998). What remains of the axoneme is then enveloped by the developing midgut and defaecated after hatching (Pitnick and Karr, 1998). Unlike the axoneme proper, the centrioles contributed by the sperm are maintained in the zygote of most metazoan species, where they function as microtubule organizing centers. Therefore, it is important that axoneme and centrioles are disconnected from one another to enable such different fates (reviewed in Schatten et al., 2011).

There are several instances in which the elimination of centrioles is also critical for proper cellular behavior (reviewed in Kalbfuss and Gönczy, 2023). This is the case for instance during oogenesis in metazoan systems, where such elimination is essential to prevent supernumerary centrioles in the zygote, since two centrioles are already contributed by the sperm in most systems. Failure to remove centrioles from the oocyte can result in multipolar spindle assembly and failed embryonic development. In *Drosophila*, centriole removal during oogenesis is modulated by the Polo kinase (Pimenta-Marques et al., 2016). Polo is present at centrioles and is critical to recruit the Peri-Centriolar Material (PCM). When Polo departs from centrioles, the PCM is lost, resulting in subsequent centriole elimination. Experimentally forcing Polo to remain at centrioles leads to PCM retention and to multipolar spindles in the ensuing zygote (Pimenta-Marques et al., 2016). The mechanism of oogenesis centriole elimination appears to be different in *C. elegans*, where Polo-like kinases and the PCM do not modulate the timing of organelle removal (Pierron et al., 2023). Instead, the microtubule-binding protein SAS-1 is important: it is the earliest known component to leave centrioles during oogenesis, and centriole elimination is premature when *sas-1* function is impaired (Pierron et al., 2023). The above findings notwithstanding, understanding of the mechanisms governing elimination of axonemes and centrioles remains incomplete. Moreover, what may have been the original mode of dismantling these elements is unclear. Investigating these processes in a basal eukaryote might help address this question.

The excavate *Naegleria gruberi* is a free-living freshwater basal eukaryote. *Naegleria gruberi* (*Naegleria* hereafter for simplicity) typically exists as an amoeba feeding on bacteria (Schardinger, 1899; reviewed in Fulton, 1993). In this life form, cells do not possess centrioles or microtubules during interphase. However, a divergent set of tubulin genes is expressed transiently upon mitosis, thus ensuring assembly of a mitotic spindle in the absence of centrioles (e.g.: Velle et al., 2022; reviewed in Fulton, 2022). Remarkably, when deprived of food, amoebae transform in a stereotyped manner into a flagellate life form within 90 min (reviewed in Fulton, 1970; Fulton, 1993). During this transformation, one centriole forms *de novo* initially, and a second centriole assembles shortly thereafter (Dingle and Fulton, 1966; Fritz-Laylin et al., 2016; Fulton and Dingle, 1967; Fulton and Dingle, 1971; reviewed in Fulton, 1993). Both centrioles then template a flagellar axoneme and organize an intracellular microtubule cytoskeleton. Now equipped with two flagella, cells readily swim away. Of particular importance for this work, most flagellated cells then spontaneously revert to the amoeboid state, and in doing so lose their flagella and their internal microtubule cytoskeleton (reviewed in Fulton, 1977b). In contrast to the extensively studied amoeboid-to-flagellate transformation, reversion from the flagellated state to the amoeboid state has been scarcely investigated. As a result, the mechanisms by which axonemes are removed and centrioles may be eliminated are not resolved. *Naegleria*’s position on the evolutionary tree of life, in one of the earliest branching lineages from the Last Eukaryotic Common Ancestor (LECA) (Rodríguez-Ezpeleta et al., 2007), makes it a promising system to offer insights into the origin of these fundamental processes. Moreover, *Naegleria gruberi* is a close relative of *Naegleria fowleri*, also known as the “brain eating amoeba” (reviewed in Güémez and García, 2021). No successful treatment is available for people infected with *Naegleria fowleri*, which is almost invariably lethal. Thus, increasing understanding of *Naegleria* biology might eventually also suggest new therapeutic avenues. In this work, we set out to investigate the flagellate-to-amoeboid reversion in *Naegleria*, with the intent of unravelling the cellular mechanisms through which the flagellar axoneme and the centrioles are eliminated during this process.

## Results

### Cytological description of reversion from flagellates to amoebae in *Naegleria*

We set out to analyze the process through which *Naegleria* reverts from a flagellate to an amoeba (Fig. 1A). In these experiments, food is removed at t=0 min to initially induce synchronous transformation from amoebae to flagellates. Thereafter, starting at t=90 min, flagellated cells begin reverting towards the amoeboid state. We found that external flagella are lost from the vast majority of cells within 2 hours during this reversion process (Fig. 1B), as previously reported (reviewed in Fulton, 1970). Moreover, analyzing the fate of centrioles during reversion using immunofluorescence with antibodies against the centriolar protein Centrin, we found that the fraction of cells harboring centrioles also diminishes within 2 hours after t=90 min (Fig. 1B). The reversion process is less synchronous than the transformation process, which presents a challenge for studying the underlying mechanisms. Previous work established that brief exposure to elevated temperature or mechanical pressure can promote reversion (Dingle and Fulton, 1966; Fulton, 1977a; Fulton and Dingle, 1967; Schardinger, 1899). Accordingly, we found that flagellated cells subjected to a 20 min heat shock at t=95 min lose their flagella more synchronously (Fig. 1B). Additionally, centriole elimination is also accelerated, although less so than flagellar loss (Fig. 1B).

**Figure 1:**
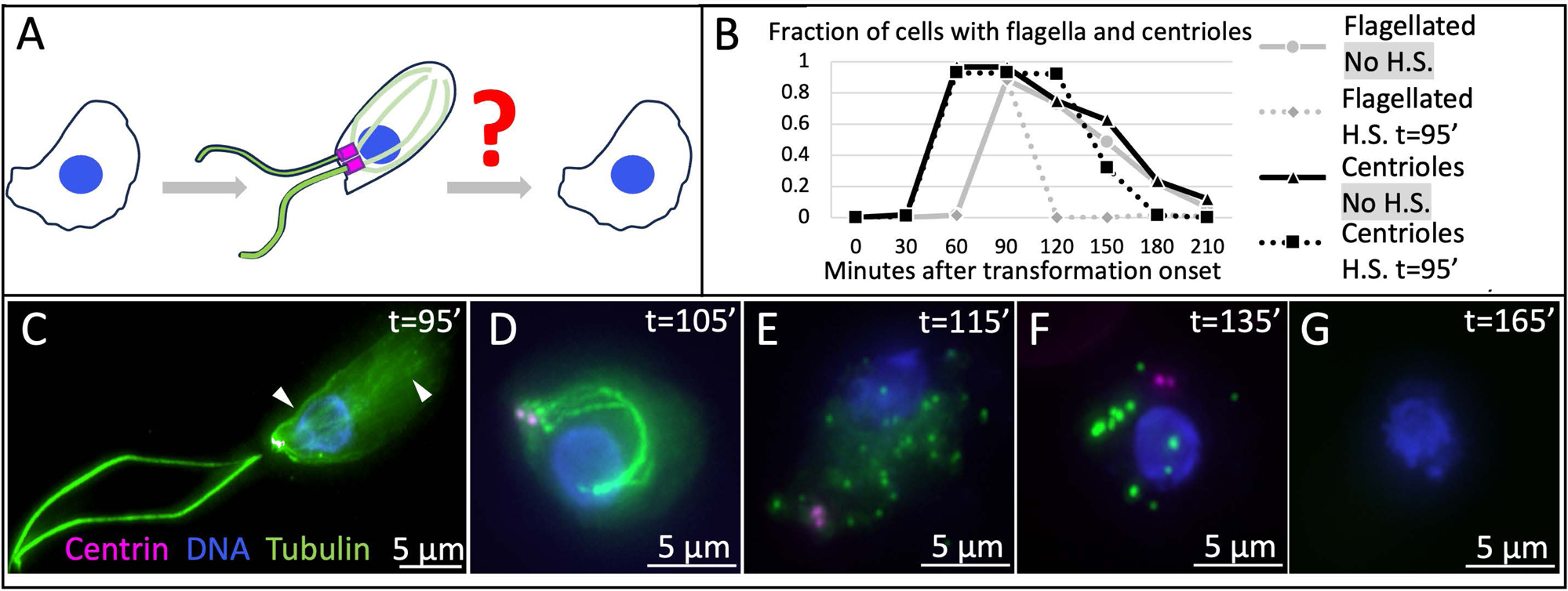
Reversion process from flagellate to amoeba in *Naegleria gruberi*. A: Schematic of reversion from flagellated life form to amoeboid life form in *Naegleria* highlighting the research question. Axonemal and intracellular microtubules: green; centrioles: magenta; nucleus: blue. B: Fraction of cells with flagella assessed after Lugol’s iodine fixation (grey), and with centrioles assessed by Centrin immunofluorescence (black). Dashed lines represent heat-shocked populations (heat shock from t=95 min until t=115 min). For each time point and condition: N > 100 cells. Unless stated otherwise, in this and all other figures, t=0 corresponds to the onset of transformation, whereas H.S. refers to heat shock. C-G: Representative images of cells undergoing reversion after heat shock (from t=95 min until t=115 min). Cells were fixed at indicated times and immunostained for α/β tubulin (green), as well as Centrin (magenta); DNA is visible in blue. Arrowheads in panel C point to intracellular microtubules.

To further investigate the elimination of flagella and centrioles, we stained cells using antibodies against tubulin to mark the axoneme and Centrin to monitor centrioles at different timepoints during reversion after heat shock. Initially, at t=95 min, we found that ∼90% of cells harbor two flagella stemming from the two centrioles located at the plasma membrane (Fig. 1C). Moreover, an intracellular microtubule cytoskeleton emanates from the two centrioles (Fig. 1C, arrowheads). At t=105 min, the flagellar axonemes are internalized in almost all cells, but remain connected to centrioles, whereas intracellular microtubules are absent, although diffuse cytoplasmic signal is still detected (Fig. 1D). Furthermore, at t=115 min most cells no longer contain flagella, but instead exhibit numerous small tubulin-positive spots, as well as diffuse cytoplasmic tubulin signal (Fig. 1E). The tubulin-positive spots probably coalesce into larger tubulin-positive elements, since at t=135 min focused tubulin signals are less numerous and brighter, and the diffuse cytoplasmic tubulin signal has been cleared (Fig. 1F; Fig. S1). Finally, at t=165 min, the majority of cells no longer exhibit signal for tubulin or centrioles (Fig. 1G). In summary, our time-resolved immunofluorescence analysis reveals the sequence of changes in axonemes and centrioles during the reversion process in *Naegleria*.

### Axonemal internalization mechanisms

We focus hereafter initially on our analysis of axoneme elimination. Intrigued by the rapid internalization of the flagellar axoneme, we conducted fast time-lapse Differential Interference Contrast (DIC) microscopy of live cells to monitor the internalization process (N=30 cells). Flagellum shedding into the extracellular milieu was never observed in these experiments. Strikingly instead, in 13/30 cells, both flagella fold back laterally onto the cell body and are internalized within milliseconds thereafter (Fig. 2A, white arrows; Movie S1). This observation is compatible with a hand drawn description reported decades ago (Rafalko, 1947). Moreover, a small bulge on the cell body is usually observed at the site of internalization immediately after contact of the flagellum (Fig. 2A, yellow arrows). In an additional 10/30 cells, this sequence of flagellar internalization could be observed for only one of the two flagella. In 2 of these 10 cases, the other flagellum was retracted inside the cell near-orthogonal to the plasma membrane over the span of several seconds (Fig. S2A). The fate of the other flagellum could not be ascertained in the other 8 cells of this subset, perhaps because it is out of focus or folds back too rapidly to be observed. Likewise, in the remaining 7/30 cells analyzed, both flagella were removed, but flagellar internalization could not be conclusively scored; in some cases, a flagellum disappeared between two frames, without any discernible indication of the path it took to presumably be absorbed in the cell.

**Figure 2:**
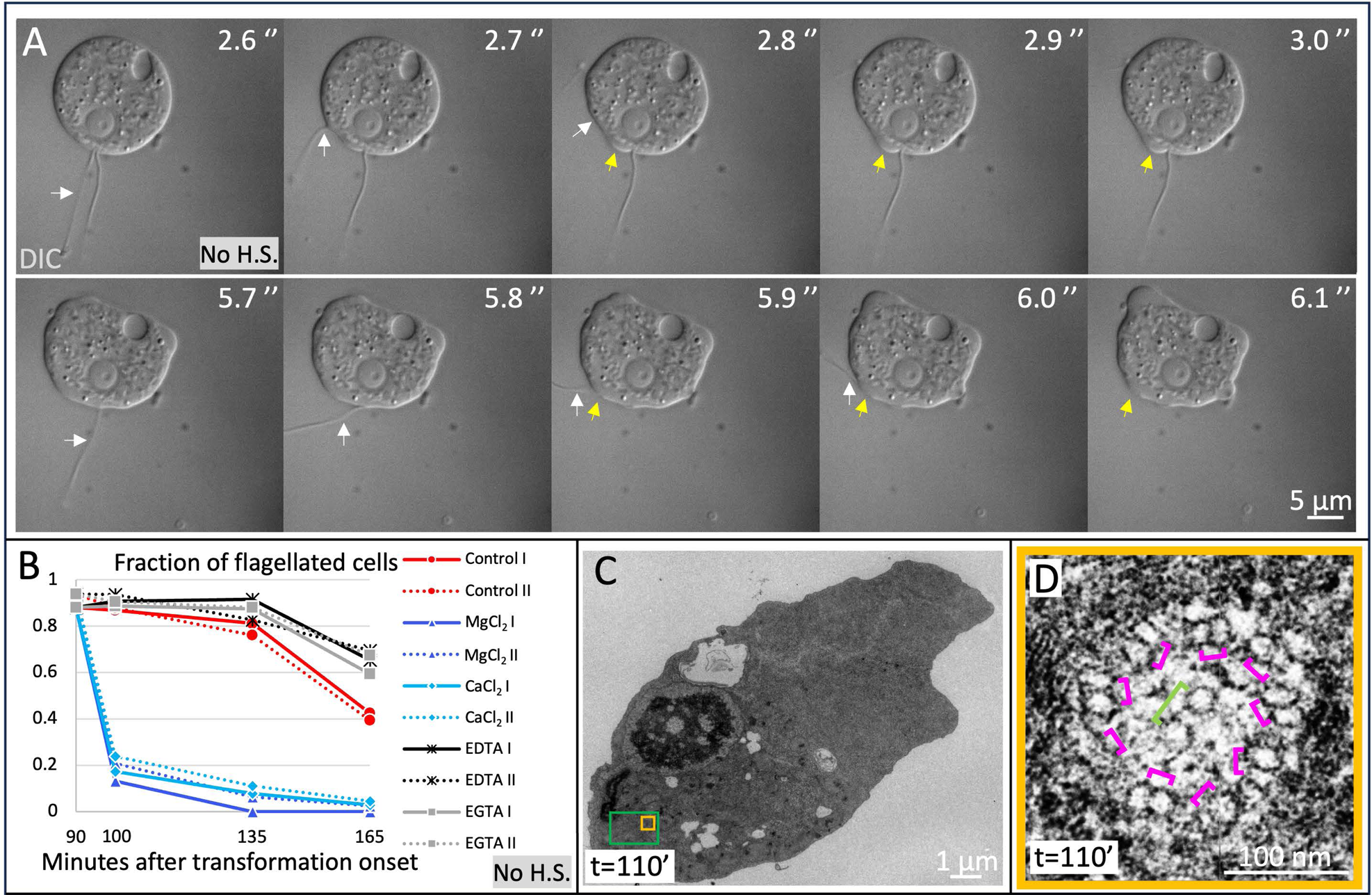
Axonemal internalization proceeds primarily through membrane fusion. A: Still images of DIC movie of flagellated cell (at approximately t=100 min). Reversal was induced by pushing down on the coverslip. Time is shown in seconds since the beginning of the movie. White arrows point to internalizing flagella, yellow arrows to membrane bulges. See also Movie S1. B: Fraction of cells with flagella assessed after Lugol’s iodine fixation. Cells were not heat-shocked. Dashed lines represent experimental repeats for each indicated condition. For each time point and condition: N > 100 cells. Note that whereas the effect of Magnesium and Calcium on transformation and reversion was tested previously (Fulton, 1977a), the ions were added at the onset of transformation in those experiments, unlike here. C: Single section EM from a serial section series, providing an overview of a cell heat shocked a t=90 min and fixed at t=110 min. Orange box indicates region magnified in panel D, green box region magnified in Fig. 3E. D: Single section EM from a serial section series, showing internalized axoneme in top view (corresponding to orange box in panel C), without membrane around the 9 microtubule doublets (indicated by magenta brackets, some better seen than others in this particular section). The pair of central microtubules is indicated by a green bracket.

We reasoned that the extremely rapid disappearance of the flagellum upon folding back laterally onto the cell body, together with the presence of a bulge thereafter, suggests that the flagellum membrane fuses with the cell body plasma membrane. To investigate this hypothesis, we treated cells with MgCl_2_ and CaCl_2_, which are both known to stimulate membrane fusion in other systems (reviewed in Mondal Roy and Sarkar, 2011). As shown in Fig. 2B, we found that addition of either MgCl_2_, as shown previously (Fulton, 1977a), or of CaCl_2_, accelerates flagellum removal, whereas addition of their chelating agents EDTA or EGTA slows down this process, together compatible with a membrane fusion mechanism. To further investigate this possibility, we conducted 50 nm-thick serial section transmission electron microscopy (EM) at t=110 min, when cells transition from having internalized axonemes to harboring small tubulin-positive spots (see Fig. 1D, 1E; N=5 cells). If axoneme internalization occurs usually via fusion between the membrane of the flagellum and that of the cell body, then internalized axonemes should not be surrounded by a membrane. Serial section EM analysis established that axonemes without a surrounding membrane are indeed present in all cells examined. That this is the case can be observed in side views (Fig. S2B), as well in rare top views in which peripheral microtubule doublets and central microtubules can be discerned (Fig. 2D; S2C). Overall, these experiments indicate that membrane fusion is the principal mechanism underlying internalization of the axoneme at the onset of the reversion process.

### Spastin-mediated axonemal fragmentation

We next turn our attention to how internalized axonemes turn into small tubulin-positive spots (see Fig. 1D, 1E). Interestingly, immunofluorescence analysis revealed that internalized axonemes exhibit stereotyped discontinuities in the tubulin signal (Fig. 3A). Thus, we found that gaps in axonemal tubulin signals are present on average every 0.26 ± 0.06 µm (SD), and exhibit a distribution distinct from that expected if fragmentation occurred at random locations (Fig. S3A). Moreover, fragmentation is observed preferentially at sites with highest axonemal curvature, and therefore highest strain (Fig. 3A, arrowheads; Fig 3B, S3B). Furthermore, analysis with STimulated Emission Depletion (STED) super-resolution microscopy revealed that the small tubulin-positive spots are circular structures with a diameter matching cross-sectional axonemal dimensions (Fig. 3C, 3D). In addition, we determined the length of axonemal fragments from side view EM images (Fig. S3C), as well as from rare top views in which consecutive serial sections were examined (Fig. 3E; Fig. S3D), revealing an average length of 0.35 ± 0.18 μm (N=6 fragments).

**Figure 3:**
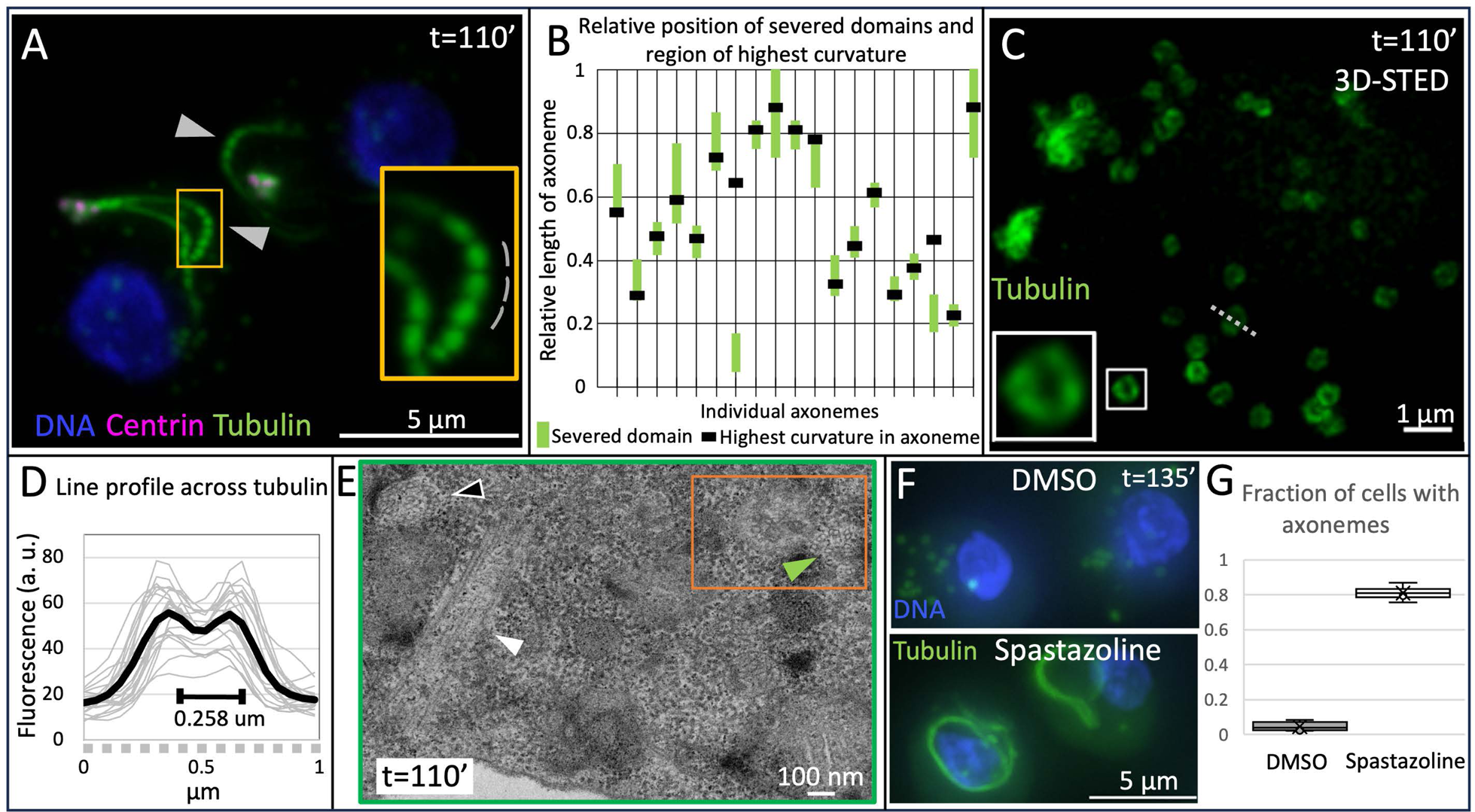
Internalized axonemes are fragmented primarily by Spastin. A: Cells heat shocked at t=95 min and fixed at t=110 min, immunostained for α/β tubulin (green), as well as Centrin (magenta); DNA is visible in blue. Arrowheads point to the most curved region of the two axonemes in the field of view. Orange box: most curved region of one axoneme, magnified on the right. The grey lines illustrate how fragment lengths were determined for Fig. S3A. B: Spatial relationship between severed domain and region of highest curvature in 19 axonemes analyzed from 2D projected 3D-stack wide-field images. Represented are the relative length of the axoneme (0: position of centriole, 1: other end of axoneme), as well as the relative position of the severed region determined by fitting a segmented line; the region with highest curvature was determined manually. Note that axonemes may still undergo movements once internalized, so that regions of maximal curvature may change over time. For the here reported data, only axonemes that displayed a single severing domain were analyzed. C: Max-projected 3D STED image of a cell heat shocked at t=95 min and fixed at t=110 min, immunostained for α/β tubulin (green). The white box highlights a circular structure that is magnified on the bottom left. The grey dashed line indicates how the line profile was determined for panel D. D: Line profiles of circular structures in panel C (grey: individual structures, black: average; N = 20 structures). E: Single section EM from a serial section series (green box in Fig. 2C). White arrowhead: internalized axonemes in side view; black arrowhead with white contour: internalized axoneme in top view, green arrowhead: axoneme fragment. Dark orange box indicates region magnified in Fig. 4A-C. F: Cells heat shocked at t=90 min for 20 min and incubated with 1% DMSO (top) or 50 μM Spastazolin (bottom); cells were then fixed at t=135 min and immunostained for α/β tubulin (green), as well as Centrin (magenta); DNA is visible in blue. G: Corresponding quantification of cells harboring an axoneme > 5 μm. N = 237 (DMSO) or 225 (Spastazolin) cells, two experimental repeats.

Which protein could promote axoneme fragmentation? We reasoned that a microtubule severing enzyme could be involved. Together with Katanin and Fidgetin, Spastin is one of three known microtubule severing enzymes containing AAA ATPase domains (Errico et al., 2002; Roll-Mecak and Vale, 2005; reviewed in Kuo and Howard, 2021), and the only one for which a specific inhibitor has been developed (Cupido et al., 2019). The *Naegleria* genome encodes a predicted Spastin homologue that exhibits high conservation within the nucleotide binding site of the AAA domain, including residues targeted by the human Spastin inhibitor Spastazoline (Fig. S3E). As shown in Figure 3F and 3G, we found that Spastazoline alters the course of axonemal fragmentation, since most cells lack axonemal fragments at t=135 min upon drug treatment. Fragments are nevertheless observed in a fraction of cells, perhaps reflecting partial inhibition of *Naegleria* Spastin by Spastazoline, or else partial redundancy with other severing mechanisms. Overall, we conclude that internalized axonemes are severed primarily by Spastin into numerous small axonemal fragments.

### Lysosome-mediated removal of axoneme-derived tubulin-containing elements

Following fragmentation by Spastin, the resulting small tubulin-containing spots presumably merge into the brighter tubulin-containing elements (see Fig. 1E and Fig. 1F), which we set out to analyze next. Interestingly, serial section EM analysis revealed the presence of membrane-enclosed elements (Fig. 4A, magenta arrowhead), which contain microtubules that are no longer organized in 9 peripheral doublets (Fig. 4A, yellow arrowhead). We found that microtubules in such membrane-enclosed elements are wider and more disorganized than those in axonemal fragments devoid of surrounding membranes (Fig. S4). Such membrane-enclosed elements could be detected in close proximity to axonemal fragments (Fig. 4B and 4C, green arrowheads). While most axonemal fragments lack membranes, as mentioned above, in rare cases a membrane could be detected around part of a fragment (Fig. 4B, green arrowhead). Given the presence of membrane-enclosed elements containing wider and more disorganized microtubules, and since the tubulin signal eventually disappears (see Fig. 1G), we hypothesized that these elements correspond to early autophagosomes, which then fuse with lysosomes, thereby resulting in tubulin removal. To test this hypothesis, we blocked lysosomal acidification using Bafilomycin A1 (Werner et al., 1984). Strikingly, this completely prevents clearance of the bright tubulin-containing elements (Fig. 4D, 4E). Moreover, this treatment also precludes clearance of the diffuse cytoplasmic tubulin signal (Fig. 4F, 4G). Interestingly, in addition, we observed that the tubulin-containing elements are not recognized by a cabazitaxel-derived microtubule probe (Fig. 4H), unlike centrioles (see below). This observation suggests that the cabazitaxel binding site is compromised in the slightly wider microtubules present in the putative autophagosomes.

**Figure 4:**
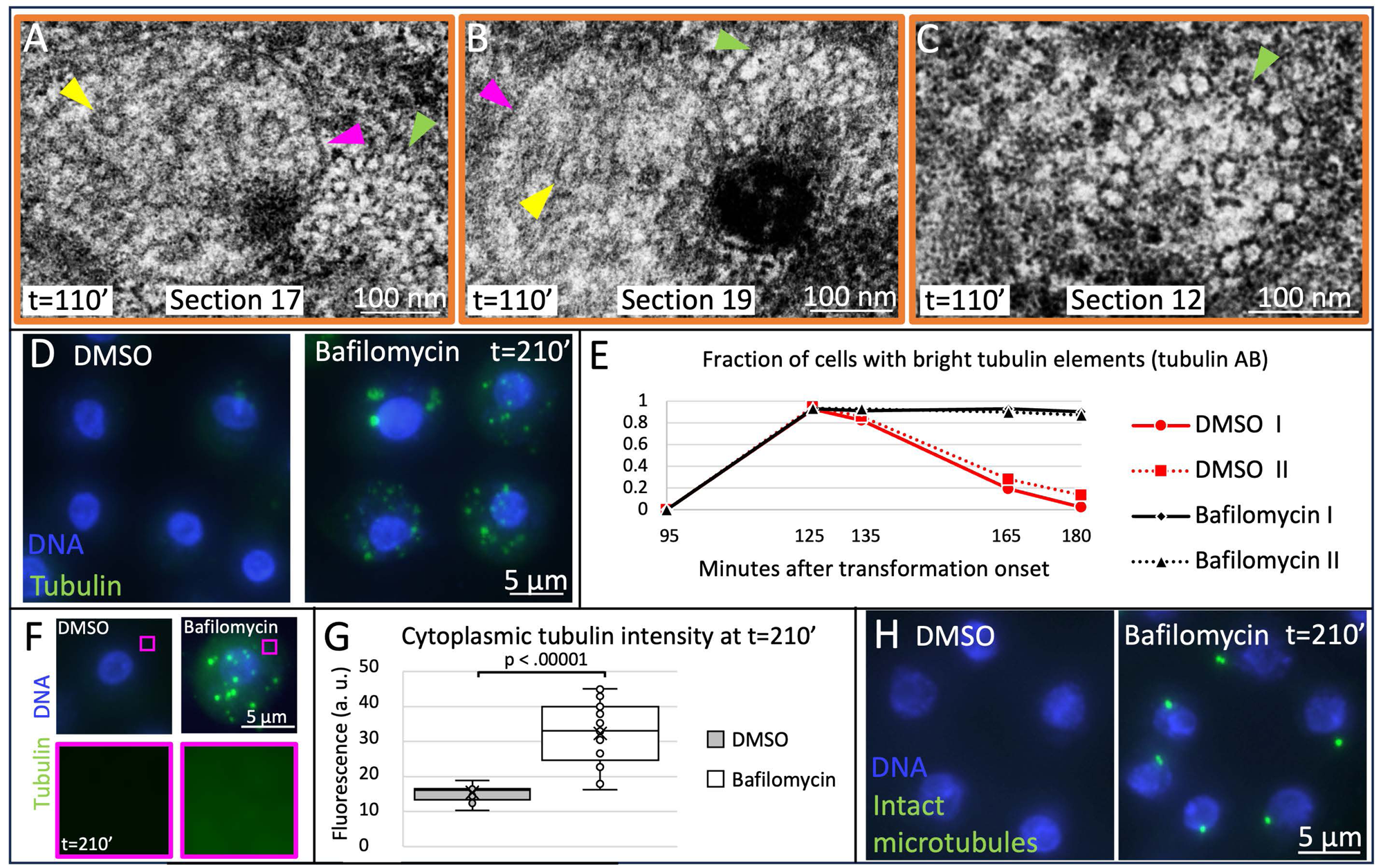
Tubulin-containing elements are membrane-enclosed and degraded by lysosomes. A-C: Single sections EM from a serial section series (dark orange box in Fig. 3E). Magenta arrowheads: membrane; yellow arrowheads: microtubule; green arrowheads: nearby axoneme fragment (partially out of focus in A and with membrane starting to enclose it in B). D: Cells heat shocked for 20 min at t=90 min and incubated with 0.1% DMSO (left) or 10 μM Bafilomycin A1 (right); cells were fixed at t=210 min and immunostained for α/β tubulin (green); DNA is visible in blue. E: Corresponding quantification of fraction of cells with bright tubulin-containing elements at indicated time points. Dashed lines represent an experimental repeat. N > 100 cells for each time point and condition. F: Same population of cells as in E. Insets highlight cytoplasmic α/β tubulin signal, which is absent in the control and present in Bafilomycin A1-treated cells. G: Corresponding quantification of cytoplasmic α/β tubulin signal intensity at t=210 min. N = 20 cells for each condition. H: Cells heat shocked for 20 min at t=90 min and incubated with 0.1% DMSO (left) or 10 μM Bafilomycin A1 (right); cells were fixed at t=210 min and immunostained with the cabazitaxel-derived probe SPY555 marking intact microtubules; DNA is visible in blue.

Overall, our findings reveal that axoneme elimination during the reversion from flagellates to amoebae in *Naegleria* follows a characteristic sequence of events. Initially, flagella fold back laterally onto the cell body and fuse with the plasma membrane within milliseconds, resulting in axoneme internalization. The axoneme is then severed in regions of highest curvature by Spastin, resulting in numerous axonemal fragments, which are then enclosed by membranes, presumably generating autophagosomes that fuse with lysosomes, ultimately leading to tubulin degradation through the lysosomal pathway.

### Centrioles are eliminated in lysosomes or in the cytoplasm, and can be shed into the extracellular milieu

We next report our analysis of centriole elimination, which occurs approximately concomitantly with disappearance of the bright tubulin-containing elements (see Fig. 1F, 1G). High spatial resolution analysis by expansion microscopy coupled to STED (U-Ex-STED) with antibodies against Centrin and tubulin revealed that, compared to intact centrioles at t=90 min, centrioles undergoing elimination have lost structural integrity at t=150 min (Fig. 5A, 5B). Moreover, we found that centrioles become wider and shorter during elimination (Fig. 5C, 5D). In addition, centrioles are progressively detached from the plasma membrane and located instead inside the cell at later stages of elimination (Fig. 5E). What mechanisms regulate centriole elimination? To begin investigating this question, we conducted live imaging of this process. Since centriole elimination occurs when cells have already resumed amoeboid motility, thereby preventing imaging of the entire elimination process within a field of view, we blocked amoeboid movements using latrunculin B to depolymerize the F-actin network (Spector et al., 1983), which did not impair axoneme or centriole elimination (see below). In these experiments, we used a cabazitaxel-derived microtubule probe (added at t=0 min) to mark centrioles, as well as Lysotracker (added at t=120 min) to mark lysosomes, given their role in removing axonemal tubulin elements in *Naegleria* (see Fig. 4D, 4E), and various cellular constituents across systems (reviewed in Trivedi et al., 2020). As shown in Fig. 5F and Movie S2, we found that in 12/25 cells in which centrioles were labelled with the microtubule probe (∼50% of all cells), centrioles are taken up by lysosomes and eliminated within them. In the remaining 13/25 cells, centrioles vanish in the cytoplasm instead (Fig. 5G and Movie S3). Strikingly, in addition, whilst conducting shorter term imaging without latrunculin B, we discovered that centrioles can also be shed into the extracellular milieu (Fig. 5H; Fig. S5A). Such externalized centrioles can be recovered by centrifugation of the culture supernatant onto a coverslip (Fig. S5B). To estimate the extent to which centriole shedding contributes to organelle removal, we filmed latrunculin B-treated cells, now adding the cabazitaxel-derived microtubule probe only at t=130 min after transformation onset. Such late addition does not allow sufficient time to label centrioles within cells, but the presence of the probe in the culture supernatant enables their detection upon shedding. This analysis established that shedding occurs in ∼10% of cells and therefore contributes in a substantial manner to centriole elimination in *Naegleria* (Fig. S5C; N=235). Moreover, since shedding also occurs in Latrunculin B-treated cells, these experiments establish that this process is actin-independent and thus not driven by exocytosis. What is the fate of centrioles shed into the external milieu? Remarkably, we discovered that externalized centrioles can be taken up by another cell (Fig. 5I; Fig. S5D-F), which is presumably followed by centriole elimination within this new host, given that no centrioles are left in the population 2 hours following heat shock (see Fig. 1B).

**Figure 5:**
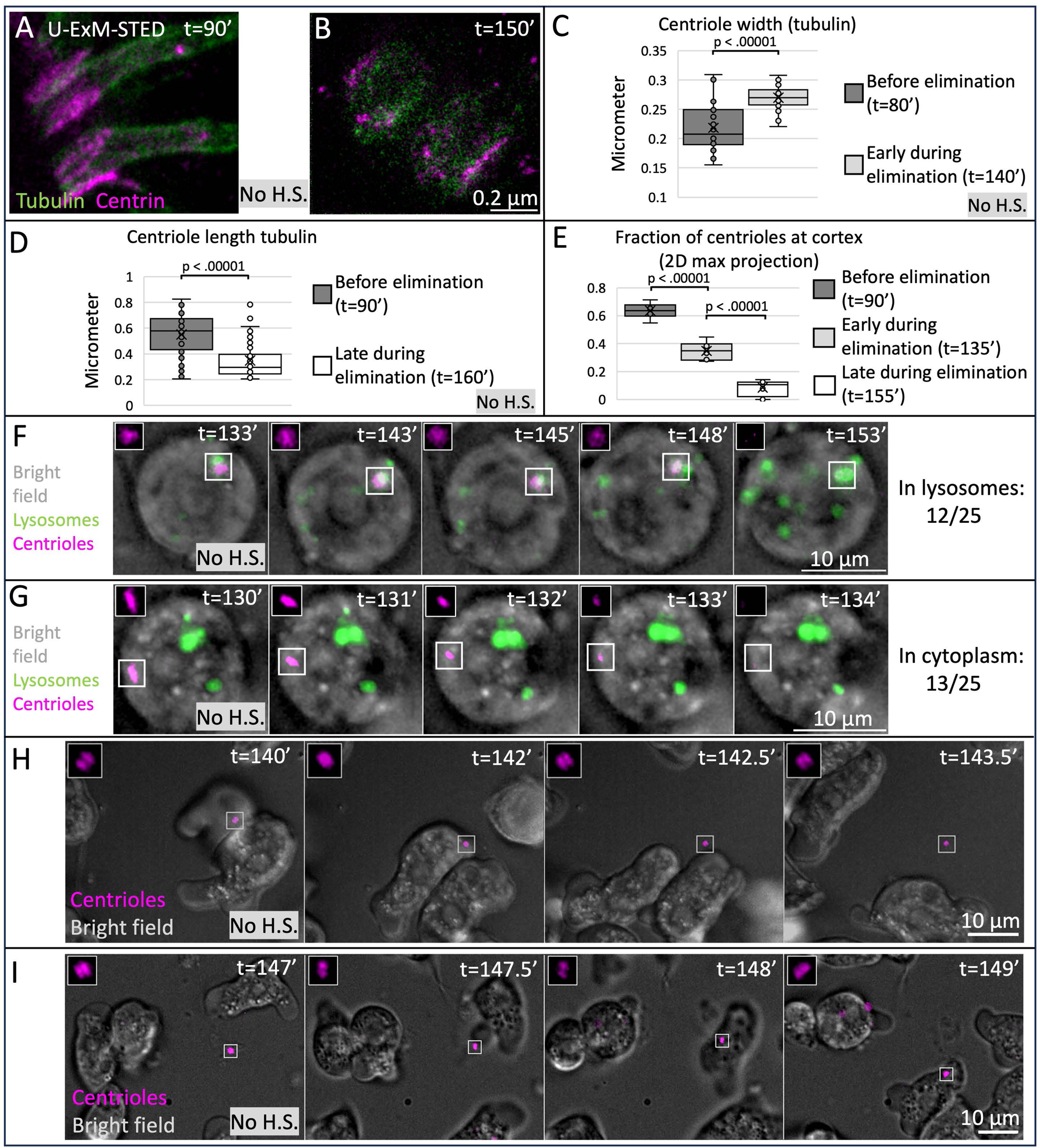
Centrioles are eliminated in lysosomes or the cytoplasm, and can be transferred between cells. A, B: Single plane U-Ex-STED images of centrioles from cells fixed at indicated time points, immunostained for α/β tubulin (green), as well as Centrin (magenta). C: Quantification of centriole width (based on α/β tubulin signal) from single-plane STED images. N=53 (t=80’) and 40 (t=140’) centrioles. D: Quantification of centriole length (based on α/β tubulin signal) from projected 3D wide-field images. N=40 centrioles for each timepoint. E: Quantification of localization of centrioles labelled by Centrin antibodies as a function of the cell border (“cortex,” visualized by brightfield and unspecific antibody background) in 2D max-projected 3D stacks of widefield microscopy images at indicated time points. Note that manual inspection of 3D images at t=90 min reveals that all centrioles are localized at the cortex, which translates into ∼60% in 2D projections. N=171 (t=90’), 190 (t=135’) and 48 (t=155’) cells. F, G: Stills of partially 2D-projected 3D stacks of widefield microscopy time-lapse at indicated time points. Amoeboid cells were immobilized with 10 μM latrunculin at the beginning of the movie, when Lysotracker was added as well; the cabazitaxel-derived microtubule probe SPY650 was present since transformation onset. Reversion to the flagellated state was induced by a 2-minute centrifugation at 600 rcf at t=95 min. F: cell in which centrioles are degraded in a lysosome G: cell in which centrioles vanish in the cytoplasm. See also Movies S2 and S3. H, I: Stills of a single plane from 3D stacks of widefield microscopy time-lapse at indicated time points. The cabazitaxel-derived microtubule probe SPY650 was present since transformation onset. Reversal to the flagellated state was induced by 2 min centrifugation at 600 rcf at t=95 min. G illustrates centriole shedding, H incorporation of that same centriole by another cell (see also Fig. S5A).

### The lysosomal pathway and the proteasome mediate centriole elimination

We next set out to test components known to modulate aspects of cellular homeostasis in other systems for their requirement in centriole elimination in *Naegleria*. Given that centrioles vanish in approximately half of the cases within lysosomes, we subjected cells to Bafilomycin A1. Strikingly, this treatment blocked centriole elimination in ∼90% of cells (Fig. 6A). Moreover, we found that the number of shed centrioles is significantly higher (Fig. S5B), presumably reflecting a shift in the balance of organelle elimination modes. Although it remains to be determined why Bafilomycin A1 affects centriole elimination in more cells than the ∼50% anticipated from colocalization experiments with Lysotracker (see Fig. 5F), these findings indicate that lysosomal-mediated removal is a major route for centriole elimination in *Naegleria*. Furthermore, given that centrioles are eliminated in approximately half of the cases outside of lysosomes, we also tested whether the proteasome is required, by treating cells with the proteasome inhibitor Bortezomib (Fissolo et al., 2008; Lioni et al., 2008). As shown in Fig. 6B, we found that Bortezomib treatment slows down centriole elimination, indicating that proteasome activity normally promotes this process. By contrast, Bortezomib has no impact on the number of shed centrioles (Fig. S5B). In line with a dual lysosome- and proteasome-mediated mechanism for centriole elimination, we found that centrioles undergoing elimination harbor autophagy-linked K63 polyubiquitination and proteasome-promoting K48 polyubiquitination (Fig 6C, 6D) (reviewed in Dósa and Csizmadia, 2022) and (Grice and Nathan, 2016)). Furthermore, we found that ubiquitin moieties are present at centrioles during the elimination process, but not before (compare Fig. 6E and 6F; Fig. 6G). Interestingly, in addition, high resolution imaging with U-Ex-STED revealed that Ubiquitin localizes together or in close proximity with microtubules (Fig. 6H), making them, or proteins closely associated with them, possible targets of ubiquitin-mediated centriole elimination.

**Figure 6:**
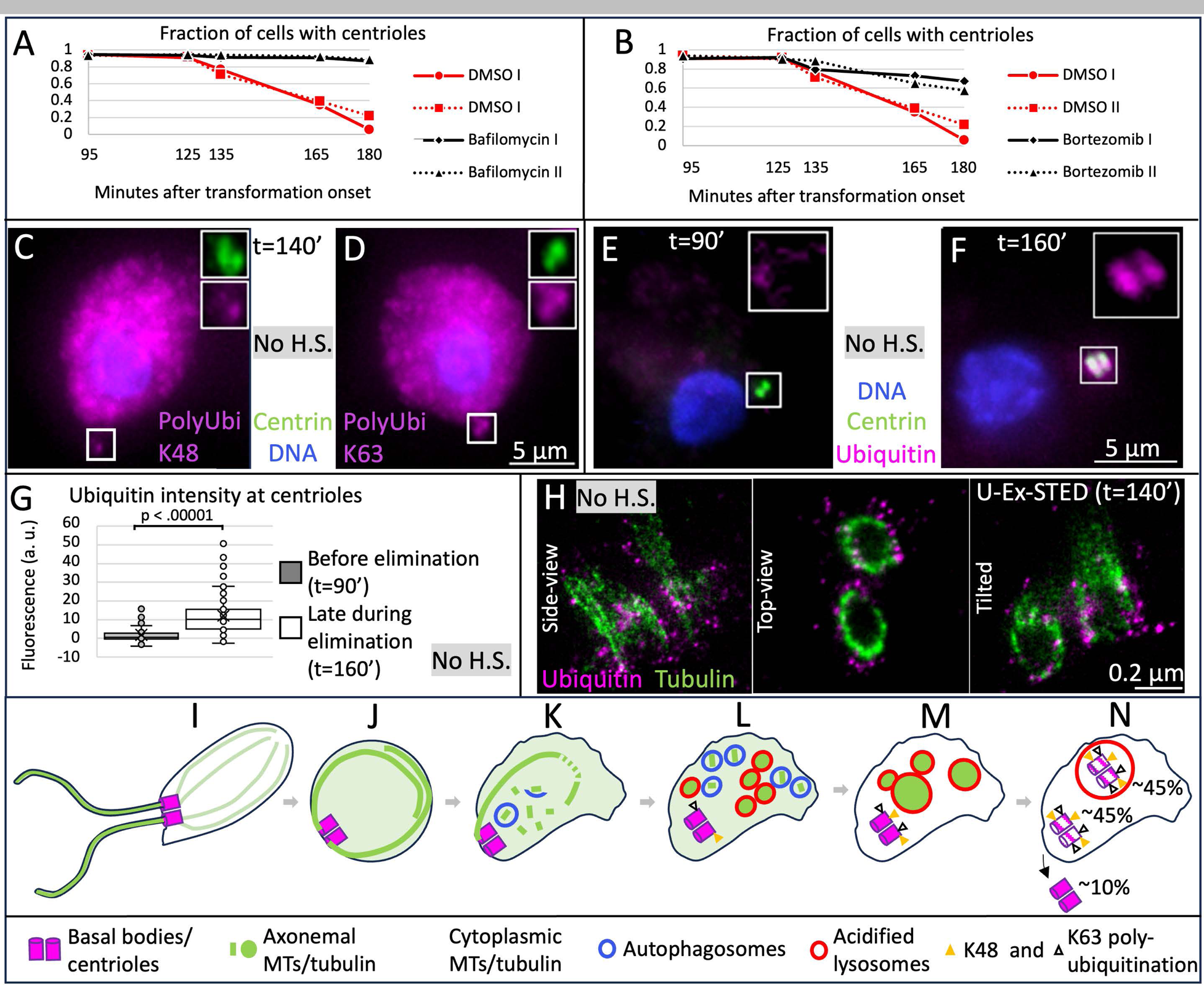
Centriole elimination depend on lysosome and proteasome function. A, B: Fraction of cells with centrioles as assessed by Centrin immunofluorescence at indicated time points; cells were heat shocked for 20 min at t=90 min. Dashed lines represent experimental repeats. Red lines: 0.1% DMSO controls; black lines: 10 μM Bafilomycin A1. C, D: Cells at t=140 min immunostained with antibodies against Centrin (green), and Poly-ubiquitin K48 (C) or Poly-ubiquitin K63 (D) (both in magenta); DNA is visible in blue. E, F: Cells at indicated time points immunostained with antibodies against Centrin (green), as well as ubiquitin (magenta); DNA is visible in blue. G: Corresponding quantification of ubiquitin signals at centrioles at indicated time points. N = 67 (t=90’) or 63 (t=160’) centriole pairs. H: Single-plane U-Ex-STED images of centrioles in indicated orientations at indicated time points. I: Flagellated cell with two centrioles and two corresponding flagella. J: Cell after internalization of axonemes following fusion of flagellar membranes with the cell membrane. K: Cell undergoing axonemal severing by Spastin at region of highest curvature, and enclosure of resulting fragments by membranes, likely early autophagosomes. L: Degradation of axonemal fragments in lysosomes. At this stage, centrioles are progressively decorated with poly-ubiquitination chains crosslinked at Lysine 48 and Lysine 63. M: Fusion of lysosomes and clearance of cytosolic tubulin. N: Elimination of centrioles either in lysosomes, the cytoplasm, or through externalization, in the indicated proportions.

Overall, our findings reveal that centriole elimination during the reversion from flagellates to amoebae in *Naegleria* entails organelle departure from the cell cortex, followed by elimination primarily in lysosomes or the cytoplasm. Such elimination is accompanied by ubiquitination in the vicinity of microtubules, with centriole removal mediated by lysosomal and proteasomal activities. In addition, centrioles can be shed into the extracellular milieu and be taken up by other cells before organelle removal.

## Discussion

The mechanisms through which axonemes and centrioles are eliminated upon reversion of *Naegleria* from flagellates to amoebae were not known prior to our work. We uncover that such elimination occurs in a step-wise fashion, as illustrated schematically in Figure 6L-Q. Flagellum removal usually entails an initial fusion event with the plasma membrane, followed by rapid internalization of the axoneme (Fig. 6M). The axoneme is then severed primarily by Spastin, yielding fragments that are subsequently enclosed by membranes and degraded through the lysosomal pathway (Fig. 6N-P). Centriole elimination entails departure from the plasma membrane, followed by poly-ubiquitination in the vicinity of microtubules, ultimately resulting in organelle removal within lysosomes or the cytoplasm (Fig. 6M-Q). Unexpectedly, we also discovered that centrioles can be shed in the external milieu and then be taken up by other cells (Fig. 6Q).

### On flagellar internalization mechanisms

We uncovered that *Naegleria* utilizes two mechanisms for flagellum removal, a main one involving rapid membrane fusion and a minor one involving slower retraction. Having two internalization mechanisms may be optimal to enable flexibility in the face of a changing environment. Moreover, it is conceivable that yet other flagellum removal mechanisms exist when *Naegleria* encounters distinct circumstances or stresses. We established that axonemes are usually internalized by membrane fusion in a low salt liquid culture, and that the pace of axoneme internalization changes by modulating bivalent cation concentration. Perhaps retention of at least two potentially ancestral internalization mechanisms reflects adaptive responses to specific environments, for instance varying salt contents or medium viscosity. Diverse modes of flagellum removal in a given organism have been reported also in evolutionary distant species (Lohret et al., 1998; Mirvis et al., 2019; Rasi et al., 2009; Venard et al., 2020; also see the review of Bloodgood, 1974). Therefore, together with our findings in the basal eukaryote *Naegleria*, it is most parsimonious to postulate that multiple mechanisms for flagellum removal co-existed early in eukaryotic evolution.

### On axoneme severing mechanisms

Following internalization, axonemes are severed with regular spacing between cut sites. Pharmacological inhibition experiments enabled us to attribute this severing activity principally to Spastin. Why do cut sites occur preferentially in regions of highest curvature? Perhaps axonemal microtubules are more readily damaged when under high strain, thereby facilitating Spastin-mediated severing. Another possibility is that Spastin activity is modulated by axonemal curvature; perhaps strain is monitored by Spastin here in an analogous manner to how tension is detected at kinetochores by the spindle assembly checkpoint (reviewed in McAinsh and Kops, 2023). For example, a negative regulator of Spastin normally present on axonemes could be removed locally as a consequence of high strain and thus allow spatially-restricted Spastin activity. Alternatively, Spastin may cut independently of strain, but severing induce the observed curvature. Regardless, axoneme fragmentation by microtubule-severing enzymes is not unique to *Naegleria*. For instance, in *Chlamydomonas*, Katanin severs doublet microtubules at the proximal end of the transition zone of the flagellum (Lohret et al., 1998; Rasi et al., 2009). Whether Spastin also plays a role in this process has not been tested. Moreover, how the single site for Katanin targeting is selected in *Chlamydomonas*, and how multiple sites undergo Spastin-mediated severing in *Naegleria* axonemes remains elusive. Furthermore, the mechanisms by which axonemal fragments are then recognized and surrounded by membranes in *Naegleria*, eventually resulting in their lysosome-mediated degradation, warrants further investigation. Unlike at centrioles, we did not detect autophagy-promoting K63 poly-ubiquitination at axonemal fragments (not shown). Perhaps other molecular marks or structural features on these fragments serve as cues for autophagosome formation and eventual lysosomal degradation. Regardless, our findings raise the possibility that axonemal fragments in other systems, for instance those derived from the sperm flagellum in the early mouse embryo (Thompson et al., 1974), are likewise engulfed by membranes before their elimination.

### Comparing centriole elimination mechanisms in fly and worm oogenesis with *Naegleria*

Comparative analyses of centriole elimination during oogenesis in flies and worms highlighted features specific the two systems (Matsuura et al., 2016; Mikeladze-Dvali et al., 2012; Pierron et al., 2023; Pimenta-Marques et al., 2016), which we discuss hereafter in the context of our findings in *Naegleria*. During *Drosophila* oogenesis, experimental retention of the Polo-like kinase 1 (Plk1) Polo at centrioles during meiotic prophase results in PCM maintenance and abrogation of centriole elimination (Pimenta-Marques et al., 2016). Could such a mechanism be at play in *Naegleria*? The genome encodes two predicted kinases with homology to Plk1 (Fritz-Laylin et al., 2010). Whether either protein is recruited to centrioles in *Naegleria* and serves a shielding role for centrioles remains to be determined. We note that although mitoses occur without centrioles in amoebae (Fulton and Dingle, 1971), in the flagellated life form centrioles seem to organize the intracellular microtubule network, which vanishes early during the reversion process, before centrioles. Therefore, it is intriguing to speculate that here also centrioles might first lose their microtubule-organizing capacity, perhaps by removal of PCM-like components, before onset of organelle elimination.

We next compare centriole elimination during *C. elegans* oogenesis and *Naegleria* reversion. Centrioles get wider during organelle elimination in both systems, although centrioles do not appear to become shorter in the worm (Pierron et al., 2023), in contrast to *Naegleria*, perhaps because worm centrioles are short cube-like structures to start with (Pelletier et al., 2006; Pierron et al., 2023; Woglar et al., 2022; Wolf et al., 1978). In *C. elegans*, the C2-containing microtubule-binding protein SAS-1 regulates the onset of centriole widening during oogenesis (Pierron et al., 2023), and perhaps a related protein exerts an analogous role in *Naegleria*. Organelle widening might be required to promote elimination through the lysosome- or proteasome-mediated pathways, conceivably by providing access for components of the ubiquitination machinery. These two elimination pathways have been investigated in *C. elegans* via downregulation of core components with RNAi, with no apparent impact on organelle demise during oogenesis (Mikeladze-Dvali et al., 2012). However, complete inhibition of these pathways blocks development, so that full chronic depletion cannot be achieved. Acute and complete depletion, for instance through a Degron-mediated system, will be required to probe their implication in the worm with certainty.

### On the regulation and importance of centriole elimination in *Naegleria*

The rapid elimination of the otherwise stable centriole during reversion in *Naegleria* suggests the existence of regulatory mechanisms that enable timely organelle removal. Besides the potential departure of PCM components, given our observation of progressive poly-ubiquitination in the vicinity of microtubules during elimination, controlled expression of E3 ligases could provide a means to initiate organelle removal. We could not identify an E3 ligase upregulated at late times after transformation onset in the existing microarray dataset (Fritz-Laylin and Cande, 2010). However, this could reflect insufficient sensitivity of the microarray experiments or post transcriptional control of E3 ligase activity. Why should *Naegleria* ensure that centrioles are removed when cells resume their amoeboid life form? In systems that use centrioles for cell division, their presence is pivotal for spindle formation and open mitosis, with supernumerary centrioles representing a threat to faithful chromosome segregation. However, as *Naegleria* undergoes vegetative growth with a closed mitotic division, in principle, the presence of two centrioles in the cytoplasm should not interfere with cell division in this case. Nevertheless, the presence of centriole in this life form may result in aberrant cellular organization and function. Preventing centriole elimination may enable to discover such potential roles, and thereby uncover the reason for which centrioles must be removed upon reversion.

In conclusion, our findings establish the detailed sequence of events that entail axoneme and centriole removal in *Naegleria*, and unveil underlying mechanisms that may operate in other systems.

## Acknowledgments

We thank Luc Reymond for providing directly labeled anti tubulin antibodies, as well as Omaya Dudin, Chandler Fulton, Elaine Lai, Marine Olivetta and Anne Villeneuve for constructive comments on the manuscript. This work was supported by grants from the Swiss National Science Foundation (310030_197749) and the European Research Council (AdG 835322) to P. G.

## Materials and Methods

### *Naegleria gruberi* culture and transformation

*Naegleria gruberi* was grown on NM plates (1 liter: 2g bacto-peptone (Difco), 11.1 mM glucose, 8.97 mM K2HPO4, 7.34 mM KH2PO4, 20g bacto-agar (Difco), autoclaved). A cell population comprising a mixture of amoebae and cysts was maintained on 10 cm diameter NM Petri dishes for 3-7 days at room temperature. For all experiments, cells were washed with water from approximately one-sixth volume of such a dish, and transferred onto a fresh 10 cm NM Petri dish, along with 500 μl of OP50(BL21) bacteria grown overnight at 37°C in LB with 50 μg/ml Ampicillin as food source. Cells were distributed evenly with a sterile spreader before incubating them overnight at 33°C. For initiating the transformation process, 10 ml of 2 mM Tris (not pH-ed) was added to the cells, which were then detached from the Petri dish with a sterile spreader, briefly spun into a pellet with a clinical centrifuge (600 rcf, 45 seconds), washed 3 times with 10 ml 2 mM Tris each, and incubated at 25.6°C under agitation (190 rpm) in a 10 ml Erlenmeyer flask. Heat shock was performed at t=90 min or t=95 min after transformation onset, leaving cells at 42°C under agitation (190 rpm) for the durations indicated in the figure legends.

### Lugol fixation for scoring of flagella

20 μl of Lugol’s iodine solution (Sigma) and 20 μl of *Naegleria* culture were mixed on a slide, and covered with a 60×24 mm coverslip, followed by immediate analysis by brightfield microscopy.

### Cell collection and fixation for immunofluorescence, U-Ex-STED, and EM

Cells were spun into a pellet with a clinical centrifuge (600 rcf, 1.5 min). For immunofluorescence and U-Ex-STED, cells were resuspended in 1 ml ice-cold (−20°C) acetone (when using SPY650 in fixed samples) or methanol (for all other immunofluorescence analyses), and fixed for 5 min on ice. This was followed by centrifugation in a table-top centrifuge (750 rcf 1.5 min) at 4°C and resuspension of the pellet in 1.5 ml PBS. Alternatively, for EM, cells were collected as above but incubated for 1 h in fixation buffer (2% Paraformaldehyde, 2.5% Glutaraldehyde in phosphate buffer 0.1M pH7.4). To mount cells on coverslips for immunofluorescence, 8 mm diameter coverslips were washed with ethanol and coated with poly-D-lysine hydrobromide (Merck, P1024); after drying, coverslips were dipped 5 times in water, then 5 times in 100% ethanol, before letting them dry again. Immediately thereafter, each coverslip was placed in a 1.5 ml Eppendorf tube containing a silicon mould. To prepare such tubes, roughly 750 μl of silicon were added, and the tubes centrifuged at 10,000 rcf for 5 minutes, resulting in the silicon reshaping so as to provide a surface perpendicular to the centripetal forces of the rotor. Such silicon-containing tubes were left to dry with an open lid for one week before use. Cells in PBS (for immunofluorescence), or in fixation buffer (for EM), were added to such Eppendorf tubes and spun in a table-top centrifuge at room temperature at 600-750 rcf for 1.5 min. Coverslips were stored until use in PBS at 4°C for immunofluorescence or washed 3 times for 5 min each in cacodylate buffer (0.1 M, pH7.4) for EM.

### Immunofluorescence and imaging

Coverslips were blocked for 20 min in 3% w/v BSA in PBS-T (PBS (pH 6.8) with 0.1% Tween-20) at room temperature. Primary antibody incubation was performed overnight at 4°C or for 2 h at room temperature in a moist chamber with primary antibodies diluted in 3% w/v BSA in PBS-T supplemented with 0.05% w/v NaN_3_. Thereafter, coverslips were washed 3 times for 5 min each in PBS-T prior to incubation with secondary antibodies for 2 h at room temperature in a moist chamber. When antibodies raised in the same species were utilized in a given experiment (for instance antibodies against Centrin and tubulin, both raised in mouse), then one component was first detected as delineated above, before re-incubating the slide with the other directly labelled primary antibody. After three 5 min washes in PBS-T, coverslips were mounted on a slide using 1.5 μl Fluoromount-G (Thermo Fisher), and hardened for 20 min at room temperature in the dark before imaging.

Antibodies used in this study: mouse anti-α-tubulin conjugated to Alexa 488 (1:500, (Lima and Cosson, 2019)), mouse anti-β-tubulin conjugated to Alexa 488 (1:500, (Lima and Cosson, 2019)), mouse anti-Centrin-2 (1: 500, Millipore Cat# 04–1624), mouse anti-Ubiquitin (P4D1, 1:500, Enzo, BML-PW0930-0100), rabbit anti-Ub-K48 (ARC0811, 1:500 Invitrogen, MA5-35382), rabbit anti-Ub-K63 (JM09-67, 1:500, Thermo Fisher, MA5-32573), goat anti-rabbit conjugated to Alexa Fluor 488 (1:1000, Thermo Fisher, A11034), as well as donkey anti-mouse conjugated to Alexa Fluor 594 secondary antibodies (1:1000, Abcam, ab150112072).

For labeling of tubulin antibodies with Alexa Fluor 488 maleimide, 3 equivalents of TCEP (from a 10 mM solution buffered to pH 7) were added at room temperature to the antibodies (typically 0.5 ml from a 1 mg/ml solution in PBS). After 1 h at room temperature, 6 volume equivalents of AF488 or AF568 fluorophore-maleimide in 10 mM DMSO solutions were added to the reduced antibody solution, and the colored mixture incubated at room temperature for 1 h. The labeled antibodies were isolated by gel filtration on a PD-10 column according to the manufacturer’s instructions, using PBS during column equilibration and elution. The labeled antibodies were stabilized with 0.01% sodium azide (1:1000 dilution from a 10% aqueous sodium azide stock solution). The degree of labeling (calculated by UV/Vis spectroscopy) was typically between 2 and 3.

Wide-field imaging was performed with a 100×/1.4 Plan-Apochromat objective on a Zeiss Axioplan 2 equipped with a motorized Z-drive (Z steps were 400 nm) and a CoolSnap ES2 camera. 2D-STED images were acquired on a Leica TCS SP8 STED 3X microscope with a 100 × 1.4 NA oil-immersion objective, using 488 nm and 589 nm excitation pulsed lasers, and 592 nm and 775 nm pulsed lasers for depletion. One-pixel Gaussian blur was applied to all images for analysis and display. For display, brightness and contrast were adjusted in the individual channels, applying the same corrections within a given series.

If not stated otherwise, quantification reported here stem from one representative experiment.

### Ultrastructure expansion microscopy

Ultrastructure expansion microscopy was performed essentially as reported (Woglar et al., 2022). Methanol-fixed coverslips were washed 3 times in PBS-T for 5 min each, followed by 2 washes in PBS for 5 min each. Coverslips were incubated overnight at room temperature in Acrylamide/Formaldehyde solution (1% Acrylamide and 1% Formaldehyde in PBS). Thereafter, coverslips were washed 3 times in PBS for 5 min each. For gelation, coverslips were incubated in 50 µl monomer solution (19% (wt/wt) Sodium Acrylate, 10% (wt/wt) Acrylamide, 0.05% (wt/wt) BIS in PBS) supplemented with 0.5% Tetramethylethylenediamine (TEMED) and 0.5% Ammonium Persulfate (APS) on a piece of Parafilm for 1 h at 37°C in a moist chamber in the dark. Coverslips with gels were incubated in denaturation buffer (200 mM SDS, 200 mM NaCl, and 50 mM Tris in distilled water, pH=9) until gels detached from the coverslips. Thereafter, gels were incubated in a 1.5 ml Eppendorf tube with denaturation buffer on a heat block at 95 °C. Gels were then transferred to 5 cm Petri dishes, washed with distilled water 5 times for 20 min each, followed by incubation in distilled water overnight at 4°C. The expansion factor was estimated by measuring the gel size with a ruler, and was typically 4-5 fold.

After expansion, gels were blocked for 1 h in blocking buffer (PBS (pH=6.8), 3% BSA, 0.1% Tween 20, sodium azide (0.05%)), followed by incubation overnight with primary antibodies diluted in blocking buffer. Gels were washed 3 times in blocking buffer for 10 min each before incubation with secondary antibodies diluted in blocking buffer (supplemented with 0.7 ug/L Hoechst (bisbenzimide H 33258)) at 37°C in the dark for 3 h. Gels were washed 3 times in blocking buffer for 10 min each before transfer into a 10 cm Petri dish for re-expansion by washing 6 times for 20 min each in distilled water. For imaging, gels were cut and mounted on a 60×24 mm coverslip coated with poly-D-lysine (Sigma, # P1024) diluted in water (2 mg/ml). A second poly-D-lysine-coated coverslip was added on top, and the specimen was sealed with hot VaLaP (1:1:1 mixture of petroleum jelly:lanolin:paraffin wax).

### Live cell imaging

For Figure 2A, Figure S2, as well as Movie S1, live cell imaging was performed using DIC microscopy of cells sandwiched between an agarose pad (3% in water) and a coverslip usually 24×60 mm in dimensions. Single planes were acquired every 100 ms. Cell movement and stage drift were corrected manually during acquisition. Brightness and contrast were adjusted for display, applying the same corrections within a given series.

For Figure 5G-I and Figure S5, inverted live dual bright-field and fluorescence microscopy was performed without a coverslip using a home-made moist chamber mounted on a 40×20 mm ethanol-washed coverslip. Imaging was performed with a 63×/1.4 Plan-Apochromat objective on a Nikon ti2 equipped with a motorized Z-drive and an Andor Zeila CMOS camera. Z-steps were 0.5 μm, with time intervals as indicated in the figure legends. All Z-steps in one channel were acquired before imaging in another channel with the same time interval. Brightness and contrast were adjusted for display; for Figure 5G and 5F and Figure S5C, a one-pixel Gaussian blur was applied.

### Electron microscopy

After washes in cacodylate buffer (see above), coverslips were fixed for 1 h with 2.5 % glutaraldehyde and 2.0 % paraformaldehyde in 0.1 M phosphate buffer (pH 7.4). Coverslips were then washed three times 5 min each in cacodylate buffer (0.1 M, pH 7.4), and then post-fixed for 40 min in the same buffer with 1.0 % osmium tetroxide and 1.5% potassium ferrocyanide, followed by 40 min in 1.0% osmium tetroxide alone. Finally, coverslips were stained for 30 min in 1% uranyl acetate in water, before being dehydrated through increasing concentrations of alcohol, and then finally embedded in epon resin (Embed 812 embedding kit, EMS). The coverslips were placed face down on glass microscope slides with a 2 mm gap between the two glass surfaces filled with resin. This gap was maintained with a 2 mm thick teflon washer. The samples were then kept in a 65°C oven for 24 h to allow the resin to harden. Finally, cells were sectioned at 50 nm thickness with a diamond knife (Diatome), and serial sections collected onto pioloform support films on single slot copper grids. These were contrasted with 2% lead citrate and 1% uranyl acetate, and images taken with a transmission electron microscope at 80 kV (Tecnai Spirit, FEI Company with an Eagle CCD camera).

### Pharmacological assays and labels

Compounds were added at the times indicated in the figure legends at the following final concentrations: MgCl_2_: 10 mM, CaCl_2_: 10 mM, EDTA: 5 mM, EGTA: 5 mM, Bafilomycin A1: 10 μM (J61835.MCR, Thermo Fisher), Latrunculin B (L5288, Merck): 10 μM, Bortezomib (10-2120, Focus): 25 μM, and Spastazoline (SML2659, Sigma): 50 μM. Lysotracker (Thermo Fisher, LysoTracker™ Green DND-26) was diluted 1:10,000 for live imaging. SPY650-Tubulin (SC503, Spirochrome) was added at 20 nM for live imaging, SPY555-Tubulin (SC203, Spirochrome) at 20 nM together with antibodies for fixed cell analysis.

### Simulating randomly positioned fragmentation events

To evaluate how distances between fragmentation events would be distributed if severing occurred at random positions along the axoneme, we simulated a random process by choosing positions on a 1D line. We defined the axoneme as a line of length L = 1.5 µm composed of 30 potential cut sites of width dx = 50 nm, reflecting experimental uncertainties in determining the exact position of a cut site. For varying the number of total cuts in the axoneme N, we randomly picked N cut sites from the 30 potential positions and computed the distances between the chosen sites as intercut distances. The distribution of expected intercut distances for N sites was obtained by collecting such distances from 1000 simulations for each N.

### Statistics

All statistical tests displayed in the figures were two-tailed Student’s t-tests performed with www.socscistatistics.com.

## Supplementary Figures and Movies legend

**Figure S1: Changes in amount and brightness of** α**/**β **tubulin spots after axoneme severing**

Quantification of number (A, N=21 cell per time point) and brightness (B, N=100 tubulin spots per time point) of α/β tubulin spots per cell at t=115 min and t=135 min after transformation onset, as indicated. Cells were heat shocked from t=95 min until t=115 min.

**Figure S2: Axoneme internalization mechanisms**

A: Still images of DIC movie of flagellated cell (∼t=100 min after transformation onset). Reversal was induced by pushing down onto the coverslip. Time in min and sec since the beginning of the movie. Arrow: retracting flagellum. Note that a bulge is visible at the end of the retracting flagellum throughout the recording. The second flagellum is present throughout the recording, but out of focus at the time of complete retraction of the first one. The second flagellum is later internalized by folding back on the cell body and subsequent membrane fusion.

B: Single section EM from a serial section series of an internalized axoneme in side view, highlighting the lack of membranes surrounding the microtubule wall at t=100 min after transformation onset. Heat shocked at t=90 min.

C: Top views of select 50 nm EM sections from 24 consecutive serial section series from an intact internalized axoneme, from the region shown in Fig. 3E. Microtubule doublets are indicated by magenta brackets, central microtubules by green arrowhead. Note that microtubules are present in all sections (i.e. >1 μm).

**Figure S3: Axonemes are severed by Spastin in regions of highest curvature**

A: Distribution of fragment length determined in the data (grey discs, binned into 0.05 μm bins; N = 98 fragments) or simulated considering random cutting (black triangles).

B: Fraction of axonemes in which the severed region coincides with the region of highest curvature. Two biological replicates (N = 74 and 52 axonemes, respectively).

C: Single section EM from a serial section series of an axonemal fragment in side view. Note that the fragment is ∼500 nm-long. Black arrowheads indicate the two ends of the fragment.

D: Top views of select 50 nm EM sections from 24 consecutive serial section series from an axonemal fragment, from the region shown in Fig. 3E. Microtubule doublets are indicated by magenta arrowheads, central microtubules by green arrowhead. Note that microtubules are present only between sections 5 and 23 (i.e. approximately 900 nm), indicative of this element being an axonemal fragment.

E: Sequence logo diagram for residues in the nucleotide-binding site of AAA domains that was used to design a specific inhibitor for Spastin (top, from: (Cupido et al., 2019)), as well as alignment of the relevant regions in human and *Naegleria* Spastin (bottom). Spastin Uniprot accession number: D2VS83 (Fritz-Laylin et al., 2010). Note that the N-terminal part of the protein was not correctly annotated and is thus missing in the accession number entry; Uniprot was contacted to correct this. Note also that mutating the “A” in “DIAGO” (highlighted in red) abolishes Spatozoline action on human Spastin (Cupido et al., 2019).

**Figure S4: Microtubule diameter in membrane enclosed vesicles and axoneme fragments**

Quantification of microtubule diameters from top views in indicated compartments. Diameters were determined by measuring the perimeter of a fitted circle in individual sections. N= 35 (A-microtubule, axonemes and axonemal fragments), 36 (central microtubules, axoneme and axonemal fragments), and 29 (microtubules, autophagosomes). Note that the diameter of the central pair of microtubules in the axoneme is larger than that of the A-microtubule in the peripheral microtubule doublets.

**Figure S5: Centrioles can be shed in the external environment and taken up by other cells**

A: All frames of the movie displayed in Figure 5H. Time is shown in min since transformation onset.

B: The supernatant of a culture at t=150 min after transformation onset, following a 20 min heat shock at t=90 min, was centrifuged onto a coverslip at 10,000 rcf for 5 minutes after removal of cells by centrifugation for 2 minutes at 750 rcf. This was followed by immunostaining for α/β tubulin (green); DNA is visible in blue. N= 6 (0.1% DMSO), 7 (25 μM Bortezomib) and 5 (10 μM Bafilomycin A1) fields of view. (right) Corresponding quantification.

C: Stills of partially 2D-projected 3D stacks of widefield microscopy time-lapse at indicated time points after transformation onset. Amoeboid cells were immobilized with 10 μM latrunculin at the beginning of the recording, when Lysotracker and the cabazitaxel-derived microtubule probe SPY650 were added. Note that in this particular case centrioles were not labelled with SPY650 to start with, and are thus not detected whilst still in the cell, but become visible upon their shedding (at t=135’). This was observed in 27/235 cells analyzed.

D: Single planes of all frames of the Z-stack movie displayed in Figure 5I. Green box indicates regions highlighted in E and F. Time in min since transformation onset.

E: Z-max projection of the region boxed in D containing the centriole.

F: 3D projection of the region boxed in D, rotated by 90° to enable viewing the cell “from the side”.

Note that until t=148.5 min, centrioles are outside the cell (cell borders are highlighted by a cyan dashed line in 3D projections -at t=148 and 148.5 min, this is visible only in the 3D projection). By contrast, starting at t=149 min, the centrioles are inside the new host.

## Supplementary Movies

**Movie S1.**

DIC movie of flagellated cell (∼t=100 min after transformation onset). Reversal was induced by pressure of the coverslip. Time in sec. Scalebar: 5 μm.

**Movies S2 and S3**

Partial Z-projection of a 3D stack of widefield microscopy time-lapse at indicated times after transformation onset. Amoeboid cells immobilized with 10 μM latrunculin at the beginning of the movie, when Lysotracker (green) was added as well; the cabazitaxel-derived microtubule probe SPY650 (magenta) was present since transformation onset. Time in min. Scalebar: 10 um.

## References

Barrett, J. T. (1912). The Development of Blastocladia strangulata, N. Sp. Botanical Gazette 54, 353–371.

Bloodgood, R. A. (1974). Resorption of organelles containing microtubules. Cytobios 9, 142–161.

Braten, T. (1971). The ultrastructure of fertilization and zygote formation in the green alga Ulva mutabilis føyn. Journal of Cell Science 9, 621–635.

Cupido, T., Pisa, R., Kelley, M. E. and Kapoor, T. M. (2019). Designing a chemical inhibitor for the AAA protein spastin using active site mutations. Nat Chem Biol 15, 444–452.

Dingle, A. D. and Fulton, C. (1966). Development of the flagellar apparatus of Naegleria. J Cell Biol 31, 43–54.

Dósa, A. and Csizmadia, T. (2022). The role of K63-linked polyubiquitin in several types of autophagy. BIOLOGIA FUTURA 73, 137–148.

Errico, A., Ballabio, A. and Rugarli, E. I. (2002). Spastin, the protein mutated in autosomal dominant hereditary spastic paraplegia, is involved in microtubule dynamics. Hum Mol Genet 11, 153–163.

Fissolo, N., Kraus, M., Reich, M., Ayturan, M., Overkleeft, H., Driessen, C. and Weissert, R. (2008). Dual inhibition of proteasomal and lysosomal proteolysis ameliorates autoimmune central nervous system inflammation. Eur J Immunol 38, 2401–2411.

Fritz-Laylin, L. K. and Cande, W. Z. (2010). Ancestral centriole and flagella proteins identified by analysis of Naegleria differentiation. J Cell Sci 123, 4024–4031.

Fritz-Laylin, L. K., Prochnik, S. E., Ginger, M. L., Dacks, J. B., Carpenter, M. L., Field, M. C., Kuo, A., Paredez, A., Chapman, J., Pham, J., et al. (2010). The Genome of Naegleria gruberi Illuminates Early Eukaryotic Versatility. Cell 140, 631–642.

Fritz-Laylin, L. K., Levy, Y. Y., Levitan, E., Chen, S., Cande, W. Z., Lai, E. Y. and Fulton, C. (2016). Rapid centriole assembly in Naegleria reveals conserved roles for both de novo and mentored assembly. Cytoskeleton (Hoboken) 73, 109–116.

Fulton, C. (1970). Amebo-flagellates as research partners: the laboratory biology of Naegleria and Tetramitus. In Methods Cell Physiol., pp. 341–476. Prescott DM, editor.

Fulton, C. (1977a). Intracellular regulation of cell shape and motility in Naegleria. First insights and a working hypothesis. J Supramol Struct 6, 13–43.

Fulton, C. (1977b). CELL DIFFERENTIATION IN NAEGLERIA GRUBERI. Annual Review of Microbiology 31, 597–627.

Fulton, C. (1993). Naegleria: A Research Partner For Cell and Developmental Biology1. Journal of Eukaryotic Microbiology 40, 520–532.

Fulton, C. (2022). The Amazing Evolutionary Complexity of Eukaryotic Tubulins: Lessons from Naegleria and the Multi-tubulin Hypothesis. Front Cell Dev Biol 10, 867374.

Fulton, C. and Dingle, A. D. (1967). Appearance of the flagellate phenotype in populations of Naegleria amebae. Dev Biol 15, 165–191.

Fulton, C. and Dingle, A. D. (1971). BASAL BODIES, BUT NOT CENTRIOLES, IN NAEGLERIA. Journal of Cell Biology 51, 826–836.

Grice, G. L. and Nathan, J. A. (2016). The recognition of ubiquitinated proteins by the proteasome. Cell Mol Life Sci 73, 3497–3506.

Güémez, A. and García, E. (2021). Primary Amoebic Meningoencephalitis by Naegleria fowleri: Pathogenesis and Treatments. Biomolecules 11, 1320.

Kalbfuss, N. and Gönczy, P. (2023). Towards understanding centriole elimination. Open Biology 13, 230222.

Kuo, Y.-W. and Howard, J. (2021). Cutting, Amplifying, and Aligning Microtubules with Severing Enzymes. Trends in Cell Biology 31, 50–61.

Lima, W. and Cosson, P. (2019). The AA344 and AA345 antibodies detect human tubulin by immunofluorescence in HeLa cells. Antibody Reports 2, e108–e108.

Lioni, M., Noma, K., Snyder, A., Klein-Szanto, A., Diehl, J. A., Rustgi, A. K., Herlyn, M. and Smalley, K. S. M. (2008). Bortezomib induces apoptosis in esophageal squamous cell carcinoma cells through activation of the p38 mitogen-activated protein kinase pathway. Mol Cancer Ther 7, 2866–2875.

Lohret, T. A., McNally, F. J. and Quarmby, L. M. (1998). A Role for Katanin-mediated Axonemal Severing during Chlamydomonas Deflagellation. Mol Biol Cell 9, 1195–1207.

Matsuura, R., Ashikawa, T., Nozaki, Y. and Kitagawa, D. (2016). LIN-41 inactivation leads to delayed centrosome elimination and abnormal chromosome behavior during female meiosis in Caenorhabditis elegans. Mol Biol Cell 27, 799–811.

McAinsh, A. D. and Kops, G. J. P. L. (2023). Principles and dynamics of spindle assembly checkpoint signalling. Nat Rev Mol Cell Biol 24, 543–559.

Mikeladze-Dvali, T., von Tobel, L., Strnad, P., Knott, G., Leonhardt, H., Schermelleh, L. and Gönczy, P. (2012). Analysis of centriole elimination during C. elegans oogenesis. Development 139, 1670–1679.

Mirvis, M., Siemers, K. A., Nelson, W. J. and Stearns, T. P. (2019). Primary cilium loss in mammalian cells occurs predominantly by whole-cilium shedding. PLOS Biology 17, e3000381.

Mondal Roy, S. and Sarkar, M. (2011). Membrane Fusion Induced by Small Molecules and Ions. Journal of Lipids 2011, e528784.

Moore, K. C. (1972). Pressure-induced regression of oral apparatus microtubules in synchronized *Tetrahymena*. Journal of Ultrastructure Research 41, 499–518.

Pelletier, L., O’Toole, E., Schwager, A., Hyman, A. A. and Müller-Reichert, T. (2006). Centriole assembly in Caenorhabditis elegans. Nature 444, 619–623.

Piao, T., Luo, M., Wang, L., Guo, Y., Li, D., Li, P., Snell, W. J. and Pan, J. (2009). A microtubule depolymerizing kinesin functions during both flagellar disassembly and flagellar assembly in Chlamydomonas. Proc Natl Acad Sci U S A 106, 4713–4718.

Pierron, M., Woglar, A., Busso, C., Jha, K., Mikeladze-Dvali, T., Croisier, M. and Gönczy, P. (2023). Centriole elimination during Caenorhabditis elegans oogenesis initiates with loss of the central tube protein SAS-1. EMBO J 42, e115076.

Pimenta-Marques, A., Bento, I., Lopes, C. a. M., Duarte, P., Jana, S. C. and Bettencourt-Dias, M. (2016). A mechanism for the elimination of the female gamete centrosome in Drosophila melanogaster. Science 353, aaf4866.

Pitnick, S. and Karr, T. L. (1998). Paternal products and by-products in Drosophila development. Proc Biol Sci 265, 821–826.

Rafalko, J. S. (1947). Cytological observations on the amoeboflagellate, Naegleria gruberi (Protozoa). Journal of Morphology 81, 1–44.

Rasi, M. Q., Parker, J. D. K., Feldman, J. L., Marshall, W. F. and Quarmby, L. M. (2009). Katanin Knockdown Supports a Role for Microtubule Severing in Release of Basal Bodies before Mitosis in Chlamydomonas. Mol Biol Cell 20, 379–388.

Ritchie, D. (1947). The formation and structure of the zoospores in Allomyces. J Elisha Mitchell Sci Soc Chapel Hill N C 63, 168–205.

Rodríguez-Ezpeleta, N., Brinkmann, H., Burger, G., Roger, A. J., Gray, M. W., Philippe, H. and Lang, B. F. (2007). Toward Resolving the Eukaryotic Tree: The Phylogenetic Positions of Jakobids and Cercozoans. Current Biology 17, 1420–1425.

Roll-Mecak, A. and Vale, R. D. (2005). The Drosophila homologue of the hereditary spastic paraplegia protein, spastin, severs and disassembles microtubules. Curr Biol 15, 650–655.

Schardinger, F. (1899). Entwicklungskreis einer Amoeba lobosa (Gymnamoeba): Amoeba Gruberi. Sitz. d. Akad. d. Wiss.(Wien), Math. Nat. Abt. 1 713–734.

Schatten, H., Rawe, V. Y. and Sun, Q.-Y. (2011). The Sperm Centrosome: Its Role and Significance in Nature and Human Assisted Reproduction. Journal of Reproductive and Stem Cell Biotechnology 2, 121–127.

Sparrow, F. K. (1937). Some Chytridiaceous Inhabitants of Submerged Insect Exuviæ. Proceedings of the American Philosophical Society 78, 23–60.

Spector, I., Shochet, N. R., Kashman, Y. and Groweiss, A. (1983). Latrunculins: novel marine toxins that disrupt microfilament organization in cultured cells. Science 219, 493–495.

Sutovsky, P. (2018). Review: Sperm–oocyte interactions and their implications for bull fertility, with emphasis on the ubiquitin–proteasome system. Animal 12, s121–s132.

Thompson, R. S., Smith, D. M. and Zamboni, L. (1974). Fertilization of Mouse Ova in Vitro: An Electron Microscopic Study*. Fertility and Sterility 25, 222–249.

Trivedi, P. C., Bartlett, J. J. and Pulinilkunnil, T. (2020). Lysosomal Biology and Function: Modern View of Cellular Debris Bin. Cells 9, 1131.

Velle, K. B., Kennard, A. S., Trupinić, M., Ivec, A., Swafford, A. J. M., Nolton, E., Rice, L. M., Tolić, I. M., Fritz-Laylin, L. K. and Wadsworth, P. (2022). *Naegleria*’s mitotic spindles are built from unique tubulins and highlight core spindle features. Current Biology 32, 1247–1261.e6.

Venard, C. M., Vasudevan, K. K. and Stearns, T. (2020). Cilium axoneme internalization and degradation in chytrid fungi. Cytoskeleton (Hoboken) 77, 365–378.

Werner, G., Hagenmaier, H., Drautz, H., Baumgartner, A. and Zähner, H. (1984). Metabolic products of microorganisms. 224. Bafilomycins, a new group of macrolide antibiotics. Production, isolation, chemical structure and biological activity. J Antibiot (Tokyo) 37, 110–117.

Williams, N. E. and Frankel, J. (1973). REGULATION OF MICROTUBULES IN TETRAHYMENA : I. Electron Microscopy of Oral Replacement. Journal of Cell Biology 56, 441–457.

Woglar, A., Pierron, M., Schneider, F. Z., Jha, K., Busso, C. and Gönczy, P. (2022). Molecular architecture of the C. elegans centriole. PLOS Biology 20, e3001784.

Wolf, N., Hirsh, D. and McIntosh, J. R. (1978). Spermatogenesis in males of the free-living nematode, Caenorhabditis elegans. J Ultrastruct Res 63, 155–169.

